# AMPKγ2 Deacetylation Drives Nuclear Translocation and Doxorubicin-Induced Cardiomyopathy via Nucleolar Stress Signaling

**DOI:** 10.64898/2026.07.16.737177

**Authors:** Canrong Li, Tiantian Yi, Yuting Cui, Baicheng Cheng, Juan Lan, Charnlo Zhang, Cha Lin, Fan Yang, Ye Chen, Xingwu Wang, Hong Peng, Bin Zhao, Leping Yan, Hongmei Tan, Xiaoduo Xie

## Abstract

Doxorubicin (Dox)-induced cardiomyopathy (DIC), characterized by cardiomyocyte apoptosis, remains a major clinical challenge in chemotherapy. The regulatory γ2 subunit of AMP-activated protein kinase (AMPKγ2) plays a key role in cardiovascular diseases, but its function in DIC is poorly understood. Here, we report that Dox induces isoform-specific deacetylation and nuclear accumulation of γ2, triggering nucleolar stress and p53-mediated apoptosis. Mechanistically, HDAC3 and TIP60 interact with γ2 and modulate the acetylation of multiple lysine residues within its nuclear localization signal (NLS), controlling its nucleocytoplasmic shuttling. Dox enhances HDAC3-mediated γ2 deacetylation, thereby driving nuclear accumulation of the γ2-containing AMPK (γ2-AMPK) while suppressing the cytosolic AMPK activity. Nuclear γ2-AMPK phosphorylates and inactivates TIF-IA, a key RNA polymerase I-specific transcription initiation factor, leading to nucleolar stress through inhibition of rRNA transcription. rRNA deficit triggers release of free ribosomal proteins (RPs), which bind to and inhibit the E3 ubiquitin ligase MDM2, resulting in p53 stabilization and activation of apoptotic signaling. Using genetically engineered cardiomyocytes and a DIC mouse model, we found that a deacetyl-mimetic γ2 mutant (6KR) exacerbated DIC, whereas an acetyl-mimetic mutant (6KQ) was cardioprotective. Collectively, our findings establish acetylation-driven nuclear translocation of γ2 as a critical node linking Dox-induced nucleolar stress to p53-dependent apoptosis and suggest a promising cardio-oncology strategy that combines HDAC inhibitors with Dox to mitigate DIC.

**Significance statement:** Doxorubicin is an effective cancer drug, but its use is limited by cardiomyopathy. Our study reveals that doxorubicin drives HDAC3-mediated deacetylation of AMPKγ2, exposing its nuclear localization signal and redirecting γ2-containing AMPK from the cytoplasm to the nucleus. Nuclear AMPKγ2 phosphorylates TIF-IA, suppresses ribosomal RNA synthesis, and activates a nucleolar stress pathway that stabilizes p53 and promotes cardiomyocyte apoptosis. In mice, a deacetylation-mimetic AMPKγ2 mutant worsens doxorubicin-induced cardiomyopathy, whereas an acetylation-mimetic mutant is protective. These findings uncover an acetylation-controlled spatial switch in AMPK signaling and identify the AMPKγ2 deacetylation–nucleolar stress axis as a potential target for reducing chemotherapy-associated cardiac injury.

## Introduction

Doxorubicin (Dox) remains a cornerstone of chemotherapy for cancer patients due to its potent anti-tumor efficacy (1, 2). Nonetheless, its clinical use is constrained by cumulative, dose-dependent cardiotoxicity that can lead to irreversible cardiomyopathy and heart failure, a condition known as doxorubicin-induced cardiomyopathy (DIC) (3–5). The molecular underpinnings of DIC are complex, involving energetic, oxidative, and genotoxic stresses that ultimately trigger typical p53-mediated apoptosis (6, 7). Dox also potently induces nucleolar stress, characterized by rRNA inhibition and the disruption of ribosome biogenesis (8–10). However, the precise molecular pathways that dictate cardiomyocyte survival or cell death remain a subject of intense investigation.

The energy sensor AMPK is a well-established master regulator of cellular energy homeostasis (11, 12). In the heart, AMPK functions as a critical stress sensor, protecting against ischemia and metabolic stress (8, 13); yet, its precise role in DIC remains poorly understood, as it can exert both pro-apoptotic and anti-apoptotic effects depending on the context and isoform involved (13, 14). AMPK exists as 12 heterotrimers composed of a catalytic α subunit (two isoforms), a regulatory β subunit (two isoforms), and an adenylate-binding γ subunit (three isoforms) (12, 15). AMPKγ senses cellular AMP/ATP level and allosterically activates the α subunit to phosphorylate downstream effectors (16, 17). Among the three γ isoforms, γ2 (encoded by *PRKAG2*) is predominantly expressed in the heart and has been implicated in both cardiac physiology and an inherited cardiomyopathy known as PRKAG2 syndrome (18, 19). Although Dox has been reported to suppress global AMPK activity, contributing to metabolic failure and cell death (20, 21), whether and how the γ2-containing AMPK (γ2-AMPK) contributes to DIC remains largely unexplored.

AMPK function is highly dependent on post-translational modifications (PTM) and subcellular localization (12, 22, 23). Extensive studies have focused on AMPK phosphorylation, particularly the essential Thr172 phosphorylation of the α subunit by LKB1 or CaMKK2 for catalytic activation (12, 24). In contrast, AMPK acetylation remains poorly characterized, with only limited regulatory mechanisms and functions identified (22, 25, 26). Furthermore, compartmentalization of AMPK complexes with specific subunit isoforms enables spatially restricted signaling, allowing the kinase to engage diverse effectors in distinct subcellular localizations, thereby ensuring context-dependent metabolic and stress responses. Accordingly, AMPK α2, β1, and γ2 subunits were reported to undergo nucleocytoplasmic shuttling, enabling differential and composition-specific functions in the cytosol and nucleus (8, 27–29). For instance, cytoplasmic AMPK phosphorylates ACC1/2 to regulate lipid metabolism and FNIP1 to regulate mitochondrial function (30–33), whereas nuclear AMPK targets epigenetic regulators and transcription factors such as TIF-IA to control rRNA transcription, ribosome biogenesis, and stress response gene expression (8, 34). Notably, γ2-AMPK emerges as a key nuclear pool in stress responses, yet the mechanisms governing its subcellular distribution remain poorly understood.

Here, we identify a novel pathway whereby Dox induces HDAC3-mediated γ2 deacetylation, leading to its nuclear accumulation. Nuclear γ2-AMPK then suppresses TIF-IA-dependent RNA polymerase I (pol I)-mediated transcription initiation, triggering nucleolar stress, ribosomal protein release, MDM2 inhibition, and p53 stabilization. Thus, γ2 deacetylation serves as a critical node linking Dox-induced nucleolar stress to cardiomyocyte apoptosis, and HDAC inhibition emerges as a potential cardioprotective strategy in DIC.

## Results

### Dox induces γ2 deacetylation and nuclear translocation

To investigate isoform-specific roles of AMPKγ subunits in DIC, we examined their subcellular localization and acetylation in response to Dox and other treatments. Unlike γ1 and γ3, γ2 contains a unique nuclear localization signal (NLS) in the extended N-terminus (Fig. 1A). AcGFP-tagged γ2, but not γ1 or γ3, translocated from the cytosol to the nucleus upon Dox or A-769662 treatment, whereas the HDAC inhibitor (HDACi) panobinostat promoted γ2 cytosolic distribution (Fig. 1B). Immunoblotting for acetyl-lysine showed that panobinostat enhanced γ2 acetylation, whereas Dox markedly reduced it; A-769662 had no effect (Fig. 1C, S1A). Notably, γ1 was distributed between the cytosol and nucleus, while γ3 (a skeletal muscle-specific isoform) was mainly cytosolic; neither responded to Dox (Fig. 1B-C). Fractionation confirmed that Dox promoted γ2 nuclear translocation, whereas panobinostat had the opposite effect (Fig. 1D). Thus, Dox specifically regulates γ2 deacetylation and nuclear localization. Using an AMPKα1/α2 double-knockout (DKO) 293T cell line (35), we found that Dox-induced γ2 nuclear accumulation occurred independently of the AMPK holoenzyme and its catalytic activity (Fig. S1B-C), implying that A-769662, which also induces γ2 nuclear translocation, likely acts through a distinct mechanism. Furthermore, panobinostat enhanced γ2 acetylation and prevented Dox-induced deacetylation and nuclear import of endogenous γ2 in multiple cell lines (Fig. 1E-F). Consistent with previous reports (20, 21, 36), Dox suppressed AMPK activity, as evidenced by reduced phosphorylation of ACC and AMPK (Fig. 1C, 1F), further confirming that AMPK activity is not required for Dox-induced γ2 translocation. Conversely, panobinostat-induced acetylation retained γ2 in the cytoplasm and sustained AMPK activity even upon Dox treatment (Fig. 1B-1F). The sirtuin inhibitor nicotinamide (NAM) had no effect on γ2 acetylation or nuclear translocation (Fig. S1D-E). Thus, Dox induces isoform-specific, acetylation-dependent cytonuclear shuttling of γ2, revealing a previously unrecognized mechanism of γ2-AMPK compartmentalization under Dox-induced stress.

**Fig. 1.**
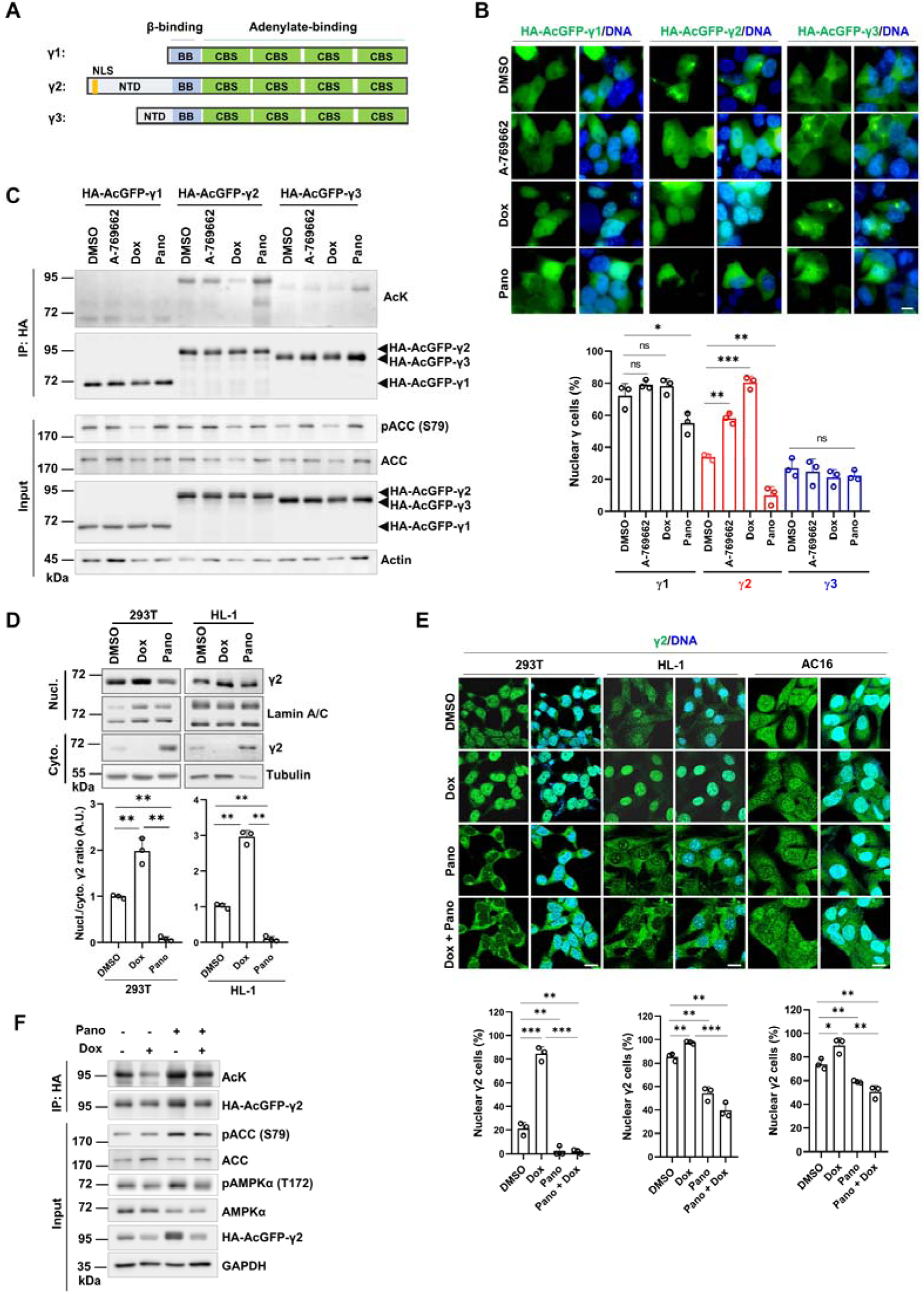
Dox induces γ2 deacetylation and nuclear translocation. **(A)** Protein structure diagram of AMPKγ subunits showing key sequence features and domain architecture. **(B)** Subcellular distribution of AcGFP-tagged γ subunits in 293T cells. Cells were transfected with γ-encoding constructs and treated with A-769662 (50 μM, 12 h), Dox (1.25 μM, 12 h), or panobinostat (Pano) (4 μM, 12 h). Representative fluorescence images (top); nuclear γ-positive cells were quantified as the percentage of cells with a nuclear/cytoplasmic signal ratio > 1.0 (n = 3, bottom). **(C)** IB analysis of γ acetylation and ACC phosphorylation. 293T cells stably expressing the three HA-AcGFP-tagged γ subunits were treated as indicated. γ acetylation was detected with an anti-acetyl-lysine (AcK) antibody after HA immunoprecipitation. **(D)** IB analysis of endogenous γ2 subcellular distribution in 293T and HL-1 cells. Cells were treated as indicated and fractionated into cytoplasmic (Cyto.) and nuclear (Nucl.) extracts. Representative blots (top). Nucl./Cyto. γ2 signal ratios were quantified and normalized to DMSO (=1) (n = 3 biological replicates, bottom). **(E)** IF staining of endogenous γ2 in 293T, HL-1, and AC16 cells. Cells were treated with Dox (1.25 μM, 12 h), Pano (4 μM, 12 h), or the combination. Representative images (top); quantified nuclear γ2-positive cells (n = 3, bottom). **(F)** IB analysis of γ2 acetylation and AMPK activity upon Dox and/or Pano treatment. 293T cells stably expressing HA-AcGFP-tagged γ2 were treated as indicated, and γ2 acetylation was detected with an AcK antibody after immunoprecipitation. Data are mean ± SD. ^✱^P < 0.05, ^✱✱^P < 0.01, ^✱✱✱^P < 0.001; ns, not significant. Statistical significance was assessed by two-tailed Student’s t-test. Scale bar, 10 μm.

### HDAC3 and TIP60 govern γ2 NLS acetylation and nucleocytoplasmic shuttling

We next explored how Dox induces γ2 deacetylation and nuclear translocation. The γ2 subunit contains a long N-terminal extension with multiple evolutionarily conserved lysines within its bipartite NLS (Fig. 1A), prompting us to examine whether reversible acetylation of these residues controls γ2 subcellular distribution. Mass spectrometry of Flag-γ2 from panobinostat-treated cells identified three acetylated lysines (K11, K12, K23) within the NLS (Fig. 2A-B). Mutating the first four (4KR) or next two (2KR) lysines to arginine only mildly reduced γ2 acetylation, whereas mutation of all six (6KR) nearly abolished it, indicating that both lysine clusters contribute to NLS acetylation (Fig. S2A). Individual K-to-R mutations revealed that each lysine contributes to deacetylation-driven nuclear translocation, with K10R and K11R showing the strongest nuclear signals (Fig. S2B). The deacetyl-mimetic 6KR mutant localized γ2 to the nucleus even under panobinostat, whereas the acetyl-mimetic 6KQ mutant (lysine-to-glutamine) retained most γ2 in the cytoplasm (Fig. 2C, S2C). To identify the responsible deacetylase (HDAC) and acetyltransferase (HAT), we performed coimmunoprecipitation (CoIP) screens. Among the tested HDACs, only HDAC3 bound γ2, and its knockout (KO) increased γ2 acetylation (Fig. 2D-E), and Dox treatment enhanced the HDAC3-γ2 interaction in cardiomyocytes (Fig. 2F). Conversely, the HAT TIP60 interacted with γ2 and promoted its acetylation: TIP60 overexpression increased, whereas KO reduced, γ2 acetylation (Fig. 2G-H, Fig. S2D-E). Immunofluorescence showed that γ2 colocalized with ectopic HDAC3 in both the cytoplasm and nucleus, with nuclear colocalization enhanced by Dox. In contrast, γ2 formed nuclear foci with overexpressed TIP60, and this colocalization was also enhanced by Dox (Fig. S2F-G). Consistently, HDAC3 KO decreased γ2 nuclear accumulation, whereas TIP60 KO increased it, phenocopying the 6KQ and 6KR mutants, respectively. Dox further promoted γ2 nuclear accumulation in WT and TIP60 KO cells, but this effect was abolished by HDAC3 KO (Fig. 2I). Furthermore, the 6KQ mutant showed impaired binding to importin α1, whereas 6KR showed enhanced binding (Fig. S2H-I), suggesting that acetylation masks the NLS and prevents nuclear import. Collectively, these results demonstrate that HDAC3 and TIP60 regulate acetylation of specific lysine clusters within the γ2 NLS, thereby controlling its nucleocytoplasmic shuttling.

**Fig. 2.**
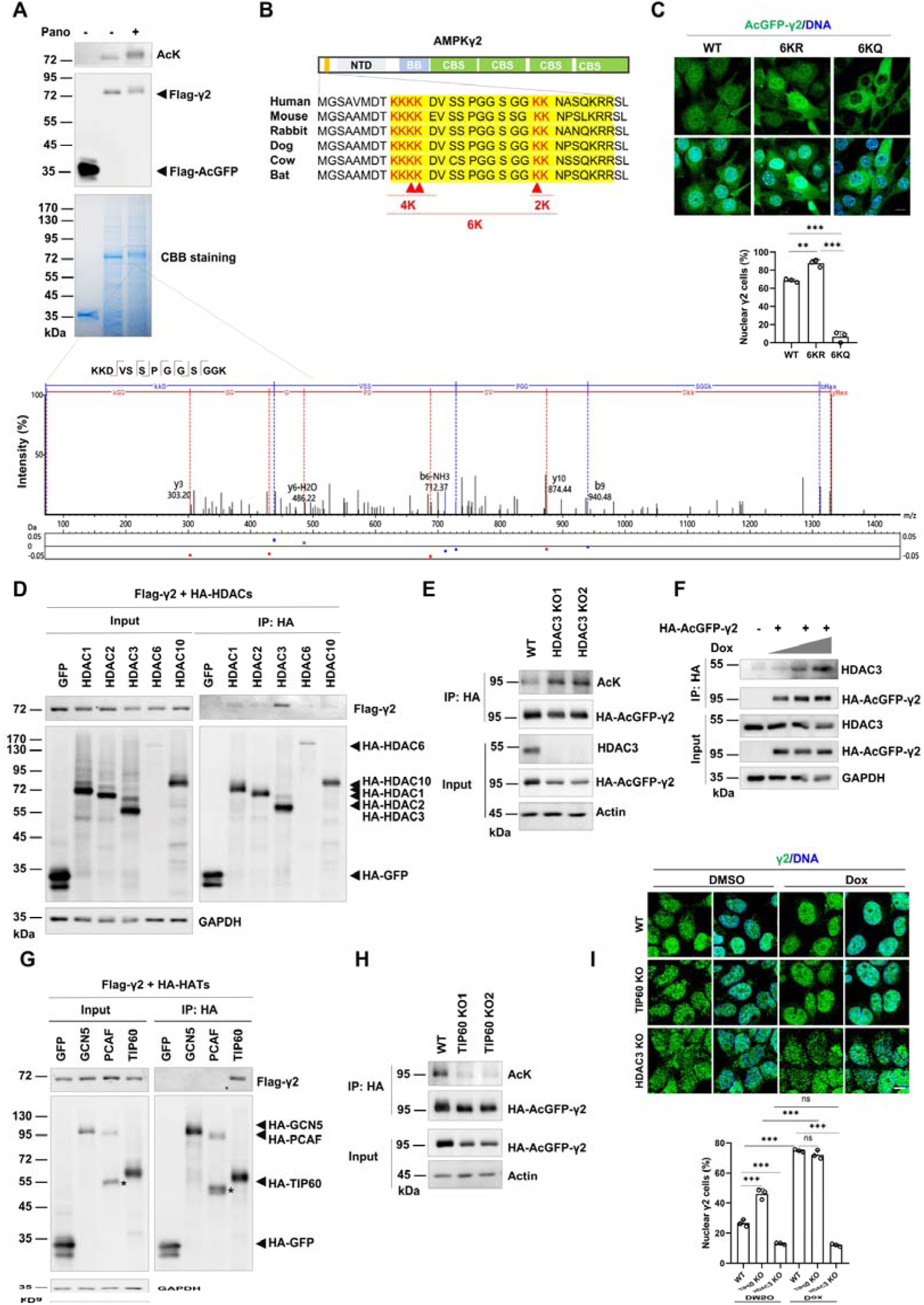
HDAC3 and TIP60 govern γ2 NLS acetylation and nucleocytoplasmic shuttling. **(A)** MS identification of acetylated lysines in the γ2 NLS upon Pano treatment. Top: IB analysis of γ2 acetylation and Coomassie blue staining of purified Flag-γ2. Bottom: MS/MS spectrum of the identified acetylated peptide. **(B)** Evolutionarily conserved lysine residues in the γ2 NLS. Sequence alignment of the γ2 NLS (highlighted in yellow) across species. Two clusters of lysine residues (in red) are conserved. K11, K12, and K23 (arrowheads) were identified by MS in (A). **(C)** Imaging of subcellular distribution of AcGFP-γ2 variants in HL-1 cells. Representative images (top); quantification of nuclear γ2-positive cells (n = 3, bottom). **(D)** CoIP screening of γ2-binding HDACs. 293T cells were cotransfected with indicated HA-tagged HDAC and Flag-γ2 constructs. Cell lysates were prepared 36 h post-transfection, and CoIP was performed using anti-HA resin. **(E)** Characterization of HDAC3 KO 293T cells and IB analysis of γ2 acetylation. **(F)** HDAC3-γ2 interaction upon Dox treatment in HL-1 cells. Cells were transfected with γ2 plasmids and treated with escalating doses of Dox (0, 1.25, 2.5 μM). The interaction was detected by CoIP. **(G)** CoIP screening of γ2-binding HATs in 293T cells, performed as in (D). **(H)** Characterization of TIP60 KO 293T cells and IB analysis of γ2 acetylation. **(I)** IF analysis of γ2 subcellular distribution in HDAC3- and TIP60-KO 293T cells. Representative images (top); quantification of nuclear γ2-positive cells (n = 3, bottom). Data are mean ± SD. ^✱^P < 0.05, ^✱✱^P < 0.01, ^✱✱✱^P < 0.001; statistical significance was assessed by two-tailed Student’s t-test. Scale bar, 10 μm.

### Dox activates nuclear γ2-AMPK and nucleolar stress signaling via TIF-IA

We next investigated the functional consequences of nuclear γ2 accumulation in Dox-induced cardiotoxicity. Dox inhibited total AMPK activity in HL-1 cells, whereas panobinostat stimulated it, as shown by AMPKα (T172) phosphorylation (Fig. S3A, S3B). Subcellular fractionation revealed that Dox activated nuclear AMPK while reducing its cytosolic activity in cardiomyocytes (Fig. 3A, S3B), indicating selective activation of the nuclear γ2-AMPK pool via γ2 nuclear translocation. To define the γ2-specific nuclear function, we generated γ1/γ2 DKO HL-1 cells rescued with γ2 acetylation variants, avoiding interference from endogenous γ isoforms (Fig. S3C). In these cells, TIF-IA, a nuclear substrate of γ2-AMPK and Pol I initiation factor, interacted more strongly with γ2 WT or 6KR than with 6KQ (Fig. 3B). Dox enhanced the γ2-TIF-IA interaction and TIF-IA phosphorylation, effects potentiated by the 6KR mutant (Fig. 3C). AMPK-mediated TIF-IA phosphorylation at Ser635 disrupts its interaction with Pol I, inhibiting pre-rRNA transcription and ribosome biogenesis (8, 34). Indeed, Dox markedly suppressed pre-rRNA synthesis in γ1/γ2 DKO cells expressing 6KR, whereas 6KQ sustained pre-rRNA levels under Dox-induced stress (Fig. 3D), indicating that γ2 deacetylation suppresses rRNA transcription via TIF-IA phosphorylation. γ2 deacetylation-induced rRNA inhibition triggered nucleolar stress in cardiomyocytes, characterized by fibrillarin (FBL) dispersion from nucleolar caps: 6KR accelerated FBL cap dispersion, whereas 6KQ suppressed it (Fig. 3E). The RP-MDM2-p53 axis is a canonical nucleolar stress pathway (37, 38). Dox-induced rRNA inhibition released free RPs, notably RPL11, which sequestered MDM2 and p53, leading to MDM2 inactivation and p53 stabilization (Fig. 3F). The 6KR mutant enhanced MDM2-RPL11-p53 complex formation and p53 stabilization more strongly than WT or 6KQ (Fig. 3G). Collectively, although Dox inhibits the major cytosolic AMPK, it induces γ2 deacetylation and nuclear accumulation, expanding the nuclear γ2-AMPK subpool. Nuclear γ2-AMPK phosphorylates and inactivates TIF-IA, causing pre-rRNA deficit, ribosome biogenesis defects, and activation of RP-MDM2-p53 nucleolar stress signaling (Fig. 3H).

**Fig. 3.**
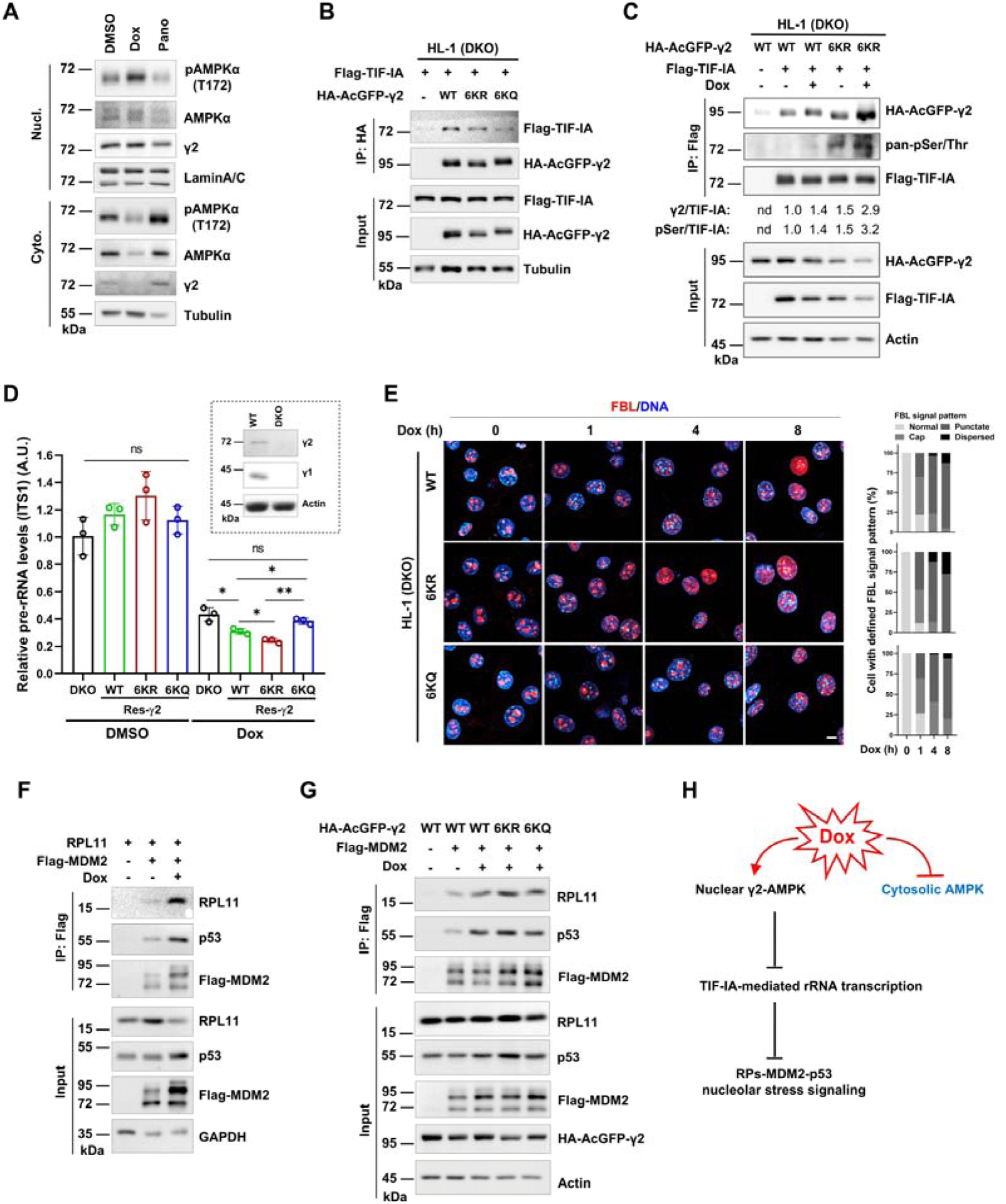
Dox activates nuclear γ2-AMPK and nucleolar stress signaling via TIF-IA. **(A)** IB analysis of AMPK activity in cytoplasmic (Cyto.) and nuclear (Nucl.) fractions upon indicated treatments. HL-1 cells were treated with Dox (1.25 μM, 12 h) or Pano (4 μM, 12 h). **(B)** Interaction between TIF-IA and γ2 variants in engineered HL-1 cell lines. γ2 variant-rescued γ1/γ2 DKO HL-1 cells stably expressing TIF-IA were subjected to CoIP using anti-HA resin. **(C)** IB analysis of TIF-IA phosphorylation and its interaction with γ2 upon Dox treatment. γ2 variant-rescued DKO HL-1 cells stably expressing TIF-IA were treated with Dox, and subjected to CoIP using anti-Flag resin. **(D)** RT-qPCR analysis of pre-rRNA levels in engineered HL-1 cell lines. γ2 variant-rescued DKO HL-1 cells were treated with Dox, and the ITS1 region of pre-rRNA was analyzed (n = 3). Inset shows γ1 and γ2 status in WT and DKO cells; γ2 rescue expression was shown in (B). **(E)** IF analysis of fibrillarin (FBL) distribution upon Dox treatment. γ2 variant-rescued DKO HL-1 cells were treated with Dox and IF-stained for FBL. Representative images (left); quantification of cells by FBL pattern (normal, cap, punctate, dispersed) (right). Scale bar, 10 μm. **(F-G)** CoIP analysis of RPL11-MDM2-p53 complex formation upon Dox treatment. **(F)** 293T cells were transfected with the indicated constructs, treated with Dox, and complexes were analyzed by CoIP using anti-Flag (MDM2); **(G)** γ2 variants and Flag-MDM2 were cotransfected, and complex formation was analyzed by Flag-MDM2 immunoprecipitation. **(H)** Schematic model illustrating the opposing effects of Dox on cytosolic AMPK (inhibition) and nuclear γ2-AMPK (activation), leading to TIF-IA inactivation and nucleolar stress signaling. Data are mean ± SD. ^✱^P < 0.05, ^✱✱^P < 0.01, ^✱✱✱^P < 0.001; ns, not significant. Statistical significance was assessed by two-tailed Student’s t-test.

### γ2 acetylation regulates Dox cardiotoxicity through p53-mediated apoptosis

To assess the functional consequence of γ2 acetylation on downstream signaling, we performed transcriptomic analysis in γ1/γ2 DKO HL-1 cells rescued with γ2 variants, with or without Dox treatment. GSEA of genotype-dependent Dox responses revealed significant enrichment of p53 pathway genes in 6KR-rescued cells compared to both WT- and 6KQ-rescued cells, whereas WT and 6KQ showed no significant difference in Dox-induced p53 transcriptional response (Fig. 4A, Supplementary Table S1). Indeed, Dox-induced p53 and cleaved caspase-3 levels were higher in 6KR cells than in WT or 6KQ cells (Fig. 4B). Cell viability assays showed that 6KR cells were more sensitive to Dox (IC_50_, 0.62 μM) than WT (IC_50_, 0.93 μM) or 6KQ cells (IC_50_, 0.93 μM), and exhibited comparable sensitivity to γ1/γ2 DKO cells (IC_50_, 0.6 μM) (Fig. 4C). Flow cytometry confirmed that 6KR cells were more susceptible to Dox-induced apoptosis (Fig. S4A). Consistently, TIP60 KO 293T cells were more sensitive to Dox, while HDAC3 KO cells were more resistant than WT cells (Fig. S4B). We further tested whether HDACi protect cardiomyocytes from Dox toxicity. Panobinostat and entinostat protected against Dox-induced cell death, whereas nicotinamide did not, and these effects were abolished in γ1/γ2 DKO cells (Fig. S4C). Drug combination synergy analysis revealed antagonism between panobinostat and Dox at higher Dox concentrations in cardiomyocytes (> 1.25 μM for HL-1 and > 2.5 μM for AC16) (Fig. 4D). Moreover, expression of the TIF-IA phospho-resistant S635A mutant counteracted Dox-induced p53 accumulation and apoptosis in both γ2 WT and 6KR cells (Fig. 4E-F, S4D), underscoring the critical role of TIF-IA phosphorylation by nuclear γ2-AMPK in DIC. Together, these data indicate that Dox-induced γ2 deacetylation promotes p53-mediated cardiomyocyte apoptosis via TIF-IA phosphorylation and nucleolar stress, and suggest panobinostat as a potential cardioprotective agent against DIC.

**Fig. 4.**
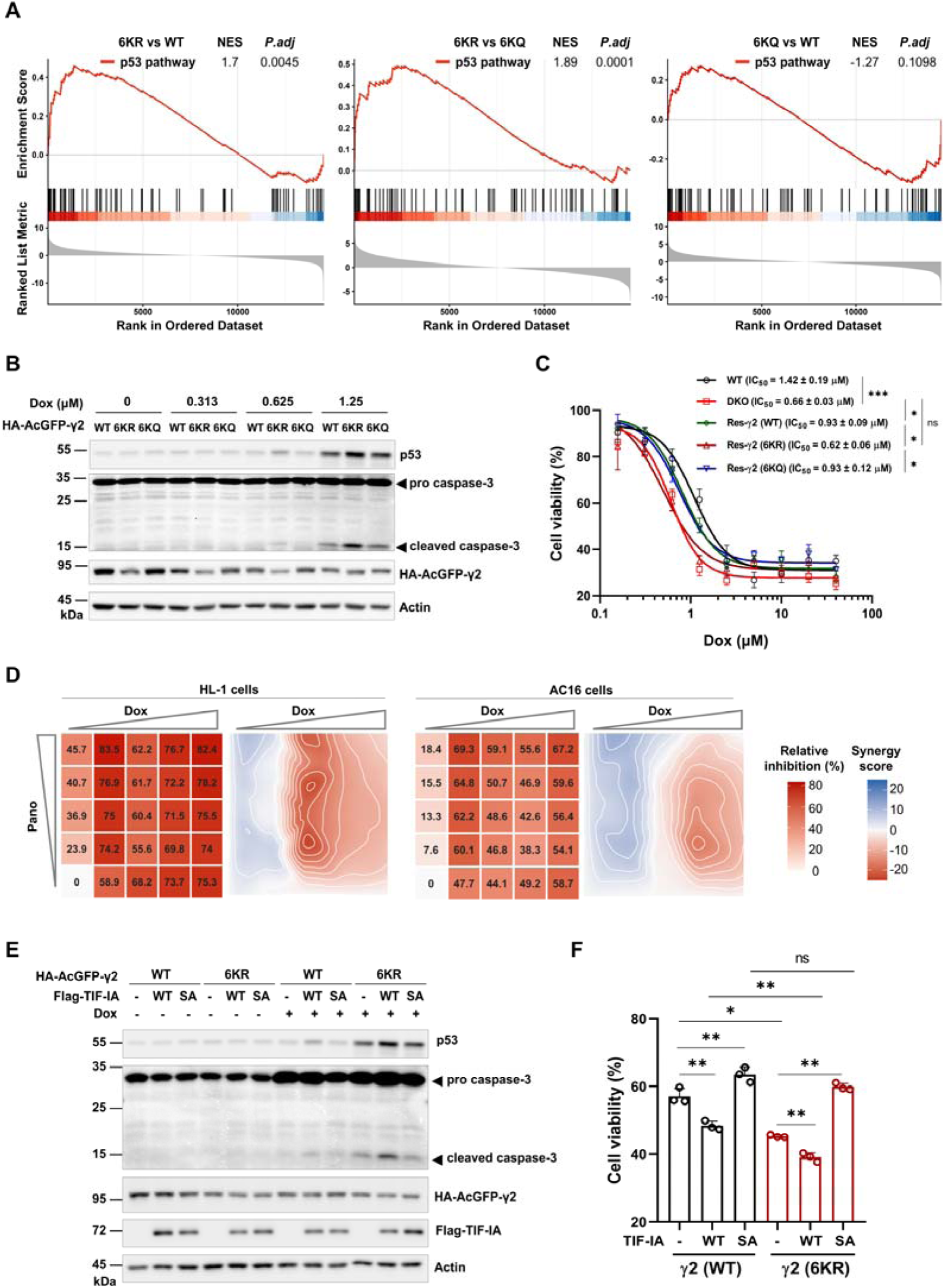
γ2 acetylation regulates Dox cardiotoxicity through p53-mediated apoptosis. **(A)** GSEA of genotype-dependent Dox responses in γ2 variant-rescued γ1/γ2 DKO HL-1 cells. The KEGG p53 signaling pathway genes enrichment in 6KR vs WT (left), 6KR vs 6KQ (middle), and 6KQ vs WT (right). P.adj, Benjamini–Hochberg–adjusted P value; NES, normalized enrichment score. **(B)** IB analysis of Dox-induced p53 and cleaved caspase-3 levels in DKO HL-1 cells expressing γ2 variants. **(C)** Dose-response curves of Dox sensitivity in DKO HL-1 cells expressing γ2 variants. Cells were treated with escalating doses of Dox (0, 0.156, 0.313, 0.625, 1.25, 2.5, 5.0, 10, 20, 40 μM, for 24 h). Cell viability was determined by CCK-8 assay. Dose-response curves were plotted, and IC_50_ values were fitted. IC_50_ values were calculated from three independent experiments and statistically compared. **(D)** Drug combination analysis in cardiomyocytes. HL-1 or AC16 cells were treated with Dox, Pano, or their combination (Dox: 0, 0.625, 1.25, 2.5, 5.0 μM for HL-1; 0, 1.25, 2.5, 5.0, 10 μM for AC16; Pano: 0, 0.08, 0.31, 1.25, 5.0 μM, for 24 h). Cell inhibition was calculated from cell viability normalized to DMSO control; heatmaps show relative inhibition across dose matrices. Contour plots display synergy scores calculated by the Bliss independence model. **(E)** IB analysis of Dox-induced p53 and cleaved caspase-3 levels in HL-1 cells. γ2 WT- or 6KR-rescued DKO cells stably expressing Flag-TIF-IA WT or S635A mutant were treated with Dox, and cell lysates were IB analyzed. **(F)** Cell viability of HL-1 cells expressing TIF-IA. γ2 WT- or 6KR-rescued DKO HL-1 cells stably expressing Flag-TIF-IA WT or S635A mutant were treated with DMSO or Dox (1.25 μM, 24 h). Cell viability was determined and normalized to DMSO control (= 100%) (n = 3). Data are mean ± SD. ^✱^P < 0.05, ^✱✱^P < 0.01, ^✱✱✱^P < 0.001; ns, not significant. Statistical significance was assessed by two-tailed Student’s t-test.

### γ2 deacetylation exacerbates DIC in mice

To validate the γ2 deacetylation-driven cardiotoxicity of Dox *in vivo*, we generated a chronic DIC mouse model using adeno-associated virus serotype 9 (AAV9)-mediated cardiac gene delivery. Mice were intravenously injected with AAV9 encoding γ2 WT, 6KR, or 6KQ under the cardiac troponin T (cTnT) promoter, followed by cumulative Dox administration (Fig. 5A). Echocardiography showed that overexpression of γ2 WT and 6KQ significantly preserved cardiac function, as indicated by higher ejection fraction and fractional shortening. In contrast, 6KR exacerbated the DIC symptoms compared to vector control (Fig. 5B). Histological analysis revealed that 6KR promoted cardiomyopathy, including chamber dilation and interstitial fibrosis (Fig. 5C-D). γ2 overexpression increases cardiac glycogen in Dox-treated mice, but with no significant difference among γ2 variants (Fig. S5A). Immunohistochemistry and immunoblot analysis confirmed elevated p53 and pro-apoptotic BAX levels in 6KR-expressing hearts following Dox treatment, with corresponding increases in TUNEL-positive apoptotic cardiomyocytes (Fig. 5E-G, S5B). In contrast, γ2 WT- and 6KQ-expressing hearts exhibited minimal p53 accumulation and reduced apoptosis (Fig. 5F, 5G), phenocopying the cardioprotective effects observed *in vitro*. These data suggest that γ2 deacetylation is sufficient to exacerbate DIC, whereas the acetyl-mimetic γ2 confers cardioprotection.

**Fig. 5.**
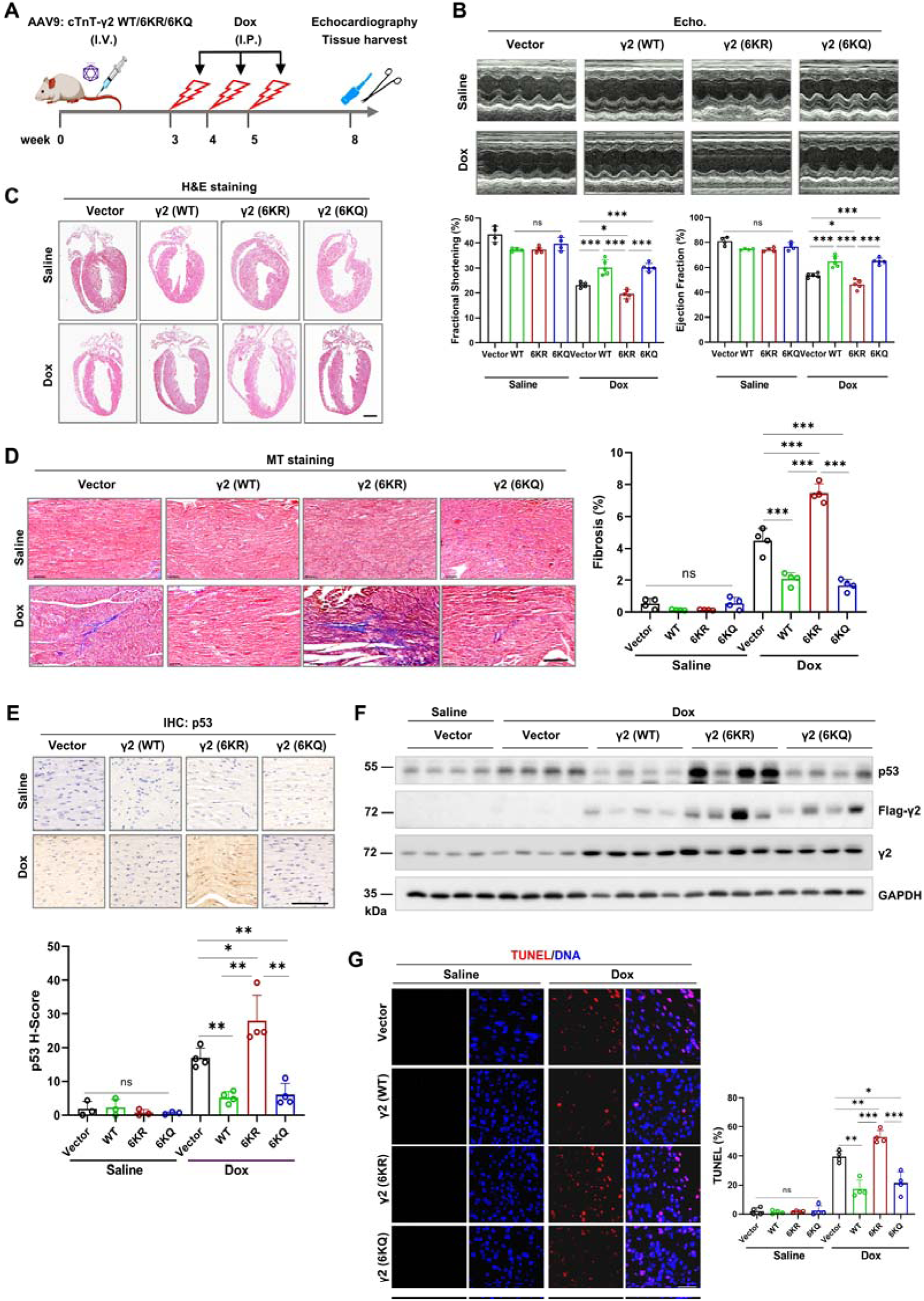
γ2 deacetylation exacerbates DIC in mice. **(A)** Schematic of the DIC mouse model protocol. Mice were injected with AAV9 encoding vector, γ2 WT, 6KR, or 6KQ intravenously, followed by cumulative Dox administration (15 mg/kg total), echocardiographic assessment, and sample preparation. **(B)** Representative M-mode echocardiography (Echo) images and quantification of fractional shortening (FS) and ejection fraction (EF) (n = 4-5 mice per group). **(C)** Representative H&E staining of heart sections. Scale bar, 1 mm. **(D)** Masson’s trichrome (MT) staining showing interstitial fibrosis (left) and quantification (right) (n = 4). Scale bar, 50 μm. **(E)** Immunohistochemical staining of p53 and H-score quantification (n = 4). Scale bar, 50 μm. **(F)** Immunoblot analysis of p53, Flag-γ2, and total γ2 expressions in DIC heart tissue. **(G)** TUNEL staining of apoptotic cardiomyocytes (red) and quantification (n = 4). Scale bar, 50 μm. Data are mean ± SD. ^✱^P < 0.05, ^✱✱^P < 0.01, ^✱✱✱^P < 0.001; ns, not significant. Statistical significance was assessed by two-tailed Student’s t-test.

Collectively, this study identifies deacetylation-driven nuclear shuttling of AMPKγ2 as a key event in doxorubicin-induced cardiomyopathy (Fig. 6). It provides a molecular framework linking isoform-specific nuclear AMPK to Dox-induced p53-dependent apoptosis in the heart and identifies panobinostat as a potential cardioprotective agent in Dox chemotherapy.

**Fig. 6.**
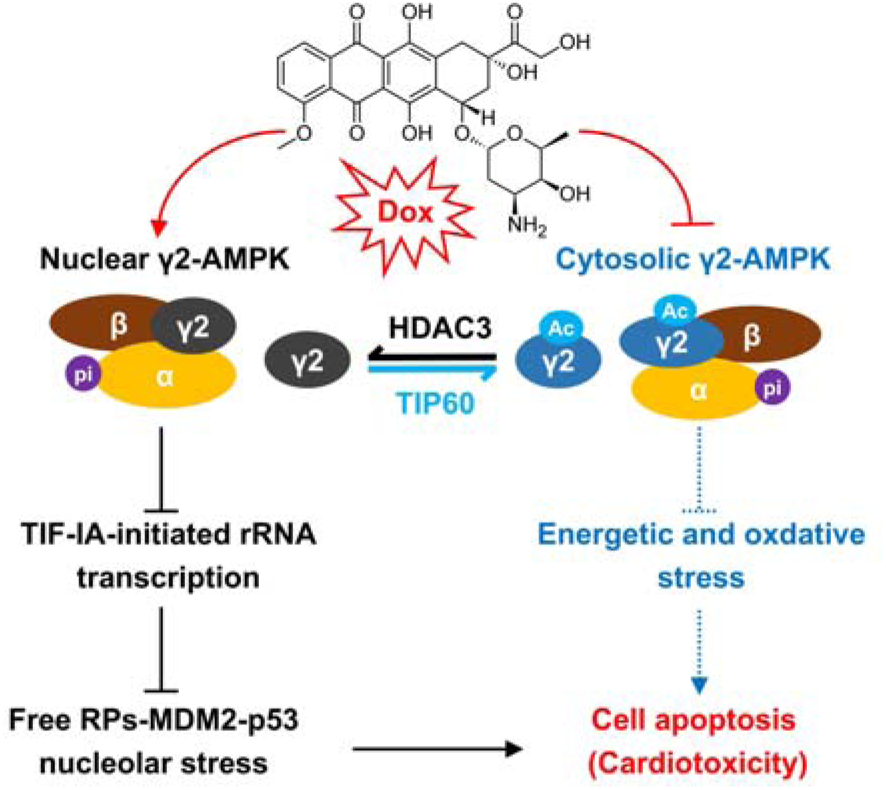
Working model of γ2 deacetylation-driven nucleolar stress in DIC. γ2 nucleocytoplasmic shuttling is dynamically regulated by TIP60-mediated acetylation and HDAC3-mediated deacetylation of its NLS. Dox promotes HDAC3 binding and deacetylation of γ2, driving nuclear γ2-AMPK accumulation and TIF-IA phosphorylation, which inhibits rRNA transcription and triggers RPs-MDM2-p53 nucleolar stress signaling. Concurrently, Dox suppresses cytosolic AMPK, exacerbating energetic and oxidative stress. Both pathways converge on p53-mediated apoptosis, leading to cardiotoxicity.

## Discussion

The cardiotoxicity of doxorubicin is a major limitation of chemotherapy. Our study reveals a unique mechanism by which Dox exploits the nucleocytoplasmic shuttling of AMPKγ2 to orchestrate a pro-apoptotic nucleolar stress response. We identified a previously unrecognized acetylation code within the γ2 NLS, dynamically regulated by TIP60 and HDAC3, that controls its subcellular localization (Fig. 2). These findings establish a mechanistic link between Dox exposure and nucleolar stress-mediated cardiomyopathy and uncover a novel isoform-specific role for γ2-AMPK in cardiac pathology.

The role of AMPK in cardiomyocyte apoptosis remains controversial, as it can exert both pro- and anti-survival effects depending on the stress context (13, 14). We found that Dox inhibits cytosolic AMPK, the major pool of total AMPK activity, consistent with previous reports (20, 21). Unexpectedly, Dox activates nuclear γ2-AMPK by promoting HDAC3-mediated deacetylation of γ2, driving its nuclear translocation and expanding the nuclear subpool. Thus, while cytosolic AMPK activation is generally protective, its nuclear accumulation in DIC is detrimental—a distinction that may explain why AMPK activators paradoxically induce apoptosis in some cell lines upon Dox treatment (39–41). Notably, the γ2-AMPK-mediated nucleolar stress pathway described here represents a p53 activation mechanism distinct from the canonical DNA damage response or AMPK direct phosphorylation of p53 (42–45). By promoting γ2 nuclear import, Dox creates a functional dichotomy: it suppresses cytosolic AMPK, exacerbating energetic and oxidative stress, while selectively activating a nuclear pool of γ2-AMPK through deacetylation. Hence, γ2 is not merely an energy sensor but also a spatial sensor of Dox-induced stresses (Fig. 6). These findings underscore that AMPK function is dictated not by global activity but by subcellular location and tissue-specific isoform composition.

TIF-IA is a known nuclear substrate of γ2-AMPK and regulates ribosome biogenesis and ER stress (8, 34). We identified TIF-IA as a key downstream effector of nuclear γ2-AMPK, linking nucleolar stress to p53-dependent cardiomyocyte apoptosis. In DIC, γ2 acts as a Dox-induced stress sensor, activating RPs-MDM2-p53 nucleolar stress signaling via TIF-IA (S635) phosphorylation and driving cardiomyocyte apoptosis (Fig. 3C-G, 4E). The convergence of nucleolar and genotoxic stress likely accounts for the robust p53 activation and potent apoptosis in DIC. In contrast to the protective role of nuclear γ2-AMPK in ischemia/reperfusion injury, where it suppresses ER stress and cell death (8), our study reveals that nuclear γ2-AMPK triggers nucleolar stress and exacerbates cardiotoxicity in DIC. Thus, γ2-AMPK mediates distinct signaling pathways and cellular outcomes in cardiac injury depending on the nature of the stress insult.

Another key discovery is the identification of HDAC3 and TIP60 as the acetylation modulators of γ2—the first such example among the three AMPKγ subunits—revealing a previously unrecognized isoform-specific regulation in the heart. HATs and HDACs are key epigenetic enzymes that control the acetylation of numerous substrates involved in cardiac pathophysiology (26, 46, 47); yet, their specific targets in DIC are largely elusive. Our work demonstrates that HDAC3-mediated deacetylation of the γ2 NLS is required for its nuclear translocation, directly linking an epigenetic modulator to nuclear γ2-AMPK regulation. However, our data cannot exclude the possibility that other HATs or HDACs (beyond HDAC3 and TIP60) may also contribute or that acetylation of residues outside the NLS could affect its nucleocytoplasmic shuttling. Given the emerging cardioprotective potential of HDACis in cardio-oncology and their synergistic anti-tumor effect with Dox (48–52), our findings have direct therapeutic implications. Specifically, the HDACi panobinostat promotes γ2 acetylation, thereby retaining γ2-AMPK in the cytoplasm. Indeed, panobinostat antagonizes Dox-induced cardiotoxicity when Dox accumulates to high concentrations (Fig. 4D, S4C). This mechanism uncouples Dox-induced stress from the p53 apoptotic axis while preserving the beneficial metabolic functions of cytosolic AMPK. Although we have not yet performed *in vivo* intervention due to the broad substrate repertoire of HDACs, panobinostat remains a promising candidate for cardio-oncology strategy in DIC.

## Material and methods

### Cell culture and drug treatment

HEK293T (293T) and AC16 cells were obtained from ATCC. HL-1 cells were purchased from Nuobo Biotechnology (Hangzhou, China). 293T AMPKα1/α2 DKO and γ1/γ2 DKO cell lines were generated as previously described (35). All cells were cultured in DMEM with 10% FBS, 100 µg/mL streptomycin, and 100 U/mL penicillin at 37 °C in 5% CO₂. Transfections were performed using PEI or Lipofectamine 3000 according to the manufacturer’s instructions. For drug treatments, compounds were added at indicated concentrations and durations as described in figure legends; control cells received equivalent DMSO or saline. For glucose starvation, cells were washed twice with PBS and cultured for 2 h in glucose-free DMEM containing dialyzed FBS (Thermo Fisher Scientific). All cell lines were confirmed mycoplasma-free before the experiment.

### Animal study

Male C57BL/6 mice (5–6 weeks old, GemPharmatech, China) were housed under specific pathogen-free (SPF) conditions and randomly assigned to experimental groups. AAV9 was delivered intravenously (10¹¹ vg/mouse). Three weeks later, one mouse per group was sacrificed to confirm cardiac γ2 expression. Dox was then administered intraperitoneally (5 mg/kg) once weekly for three weeks (cumulative 15 mg/kg) to induce DIC. Three weeks after the final Dox injection, cardiac function was assessed by echocardiography, and heart tissues were harvested for biochemical and histological analyses. All animal procedures complied with ethical guidelines and were approved by the Institutional Animal Care and Use Committee (IACUC) of Sun Yat-sen University.

### Chemical reagents, antibodies, and plasmid constructs

Dox, A-769662, panobinostat, entinostat, and nicotinamide were from TargetMol (Shanghai, China). Antibodies are listed in Supplementary Table S2. The piggyBac vectors pAir and PB were generated as previously described (53). The lentiCRISPRv2 plasmid was from laboratory stocks. pAAV-cTNT-Cre, MDM2, and TIF-IA plasmids were from Wuhan Miaoling Biotechnology. cDNAs encoding human AMPKγ1, γ2, γ3, HDAC1–3, HDAC6, HDAC8, HDAC10, GCN5, PCAF, and TIP60 were amplified from the HEK293T cDNA library and subcloned into pAir or pAAV vectors. Point mutations in γ2 and TIF-IA were generated using the Agilent QuikChange site-directed mutagenesis kit according to the manufacturer’s instructions. AAV9 constructs were generated by replacing the Cre cassette in pAAV-cTnT-Cre with Flag-γ2 (WT, 6KR, or 6KQ). All constructs were verified by Sanger sequencing. Primer and oligo sequences are listed in Supplementary Table S3.

### Immunoblot, immunoprecipitation, and immunofluorescence staining

These assays were performed as described (54). For immunoblotting, cells or tissues were lysed in buffer containing protease, phosphatase, and HDACi (panobinostat, TSA, NAM). For immunoprecipitation, 1 mg lysate was precleared with protein A/G beads, then incubated with anti-Flag or anti-HA beads for 2 h to overnight at 4 °C. Beads were washed and bound proteins were eluted in 1× Laemmli buffer by boiling at 95 °C for 15 min. For immunofluorescence, cells on polylysine-coated coverslips were treated, fixed with 4% paraformaldehyde, permeabilized with 0.1% Triton X-100, and blocked with 5% BSA. After primary antibody incubation, cells were stained with Alexa Fluor 488/647 secondary antibodies. Nuclei were counterstained with Hoechst 33342 (1 µg/mL) for 10 min, then coverslips were mounted and stored at 4 °C.

### IP-Mass spectrometry identification of acetylation sites

For acetylation site identification, ∼10⁷ HEK293T cells stably expressing Flag-γ2 were collected per sample after indicated treatments. Flag-γ2 was immunoprecipitated with 50 μL anti-Flag beads. Immunoprecipitants were resolved by SDS-PAGE and stained with Coomassie dye. The corresponding band was excised and subjected to in-gel tryptic digestion. Extracted peptides were desalted with C18 material and analyzed on an Easy-nLC 1200 coupled to a timsTOF HT (Bruker) in DDA-PASEF mode by Wininnovate Bio (Shenzhen, China). MS/MS data were searched against a human protein database with carbamidomethylation of cysteine as a fixed modification and oxidation of methionine and lysine acetylation as variable modifications. Acetylation sites were assigned based on MS/MS spectra supporting site localization.

### Virus production and generation of genetically engineered stable cell lines

For AAV9 production, HEK293T cells were cotransfected with pAAV helper, pAAV9 package, and pAAV-cTnT-Flag-AMPKγ2 constructs or vector using PEI. Viral particles were harvested at 72 h and purified by iodixanol gradient ultracentrifugation as previously described (55). Stable cell lines were generated using the piggyBac transposon system. Cells were cotransfected with a donor plasmid carrying the target gene and a transposase plasmid (5:1 ratio) using Lipofectamine 3000. After 72 h, cells were selected with hygromycin or blasticidin for 1 week to obtain pooled stable cells. For CRISPR/Cas9 gene editing, sgRNAs targeting human HDAC3, TIP60, or murine γ1 and γ2 were designed using the TrueDesign Genome Editor (Invitrogen). Oligonucleotides were cloned into lentiCRISPRv2. CRISPR/Cas9 gene editing was carried out following established protocols (56). Primer and sgRNA sequences are listed in Table S3.

### RNA sequencing and RT‒qPCR

The RNA sequencing was performed by Beijing Tsingke Biotech Co., Ltd. Raw count matrices were imported into R (version 4.2.1) and analyzed using DESeq2 (version 1.36.0). Samples were grouped by genotype (HL-1 γ1/γ2 DKO rescued with γ2-WT, 6KR, or 6KQ) and treatment (DMSO or Dox). Differential expression was modeled using a two-factor design with an interaction term, specified as ∼ cell * treatment in DESeq2, after filtering low-count genes. Genotype-dependent Dox responses were evaluated by comparing Dox-induced transcriptional changes among WT-, 6KR-, and 6KQ-rescued cells. Gene set enrichment analysis (GSEA) was performed using clusterProfiler (version 4.4.4) with genes ranked by the Wald statistic. The mouse KEGG p53 signaling pathway was obtained from the KEGG pathway database and used as the reference gene set. GSEA results are reported as normalized enrichment scores (NES) and Benjamini-Hochberg-adjusted P values; the enriched genes in each group were listed in Supplementary Table S1. For RT-qPCR, total RNA was extracted with TRIzol. Reverse transcription used 1 μg RNA, following the affiliated protocol in the StarScript III RT Kit (GenStar, China). qPCR was performed with 2× RealStar Fast SYBR qPCR Mix (GenStar). Relative RNA levels were calculated by 2^-ΔΔCt^ and normalized to Ubc. Each experiment had three technical replicates and was repeated twice. Primers are listed in Table S3.

### Subcellular fractionation

Subcellular fractionation was performed as previously described (55). Briefly, cells were lysed in low-salt buffer (10 mM HEPES pH 7.5-8.0, 10 mM KCl, 0.1 mM EDTA) with PMSF and protease/phosphatase/HDACi (panobinostat, TSA, NAM) to obtain cytoplasmic fractions. Nuclear pellets were washed and extracted with high-salt buffer (20 mM HEPES pH 7.5-8.0, 400 mM NaCl, 1 mM EDTA, 1.5 mM MgCl₂, 0.1% SDS) with the same inhibitors. Protein concentration was determined by BCA assay before immunoblotting.

### Cell viability and drug synergy

Cells were seeded in 96-well plates at densities of 4,000-10,000 cells/well (depending on cell line, 3-5 replicates). After 24 h, cells were treated with the indicated concentrations of Dox. Viability was measured by the CCK-8 assay kit (TargetMol, China) according to the manufacturer’s instructions. Viability was normalized to DMSO control. Dose-response curves were plotted using GraphPad Prism 9.0; the half-maximal inhibitory concentration IC_50_ values were calculated by nonlinear regression with a four-parameter logistic model. For drug combination synergy analysis, cells were treated with Dox and panobinostat alone or in combination; viability was assessed by CCK-8 assay, drug synergy scores were calculated in R (version 4.2.1) using the Bliss model implemented in the synergyfinder package (version 3.4.5) (57), and the results were visualized using ggplot2.

### Flow cytometry for apoptosis

Apoptosis was measured using the Annexin V YSFluor™ 647/PI kit (Yeasen, China, #40304ES60). After treatment, floating and attached cells were collected, pooled, washed twice with cold PBS, and resuspended in binding buffer. Cells were stained with Annexin V and PI for 15 min at room temperature in the dark, then analyzed on a CytoFLEX flow cytometer (Beckman Coulter). Total apoptosis was defined as the percentage of early apoptotic Annexin V^+/^PI^-^ cells plus late apoptotic Annexin V^+^/PI^+^ cells.

### Echocardiography

Cardiac function was assessed by transthoracic echocardiography using a Mindray ultrasound system (S7 SCI) as previously described (58). Briefly, mice were anesthetized with isoflurane (3% induction, 1% maintenance) and placed on a heating pad during examination. M-mode images of the left ventricle were acquired, and ejection fraction (LVEF) and fractional shortening (FS) were calculated according to the manufacturer’s instructions.

### Histological staining and immunohistochemistry

Heart tissues were fixed in 4% paraformaldehyde, paraffin-embedded, and sectioned (5 μm). Sections were stained with hematoxylin and eosin (H&E), Masson’s trichrome (MT), immunohistochemistry, TUNEL, or periodic acid–Schiff (PAS) using commercial kits. Masson’s trichrome assessed fibrosis, immunohistochemistry used indicated antibodies, TUNEL detected apoptotic cells, and PAS evaluated glycogen accumulation. Images were acquired with a KF-FL-120 digital slide scanner (KFBIO, Ningbo, China) and quantified with ImageJ.

### Fluorescence microscopy and image processing

Fluorescence images were acquired using a Nikon ECLIPSE Ti2 or Zeiss LSM 900 confocal microscope (60× oil objective) and processed with ImageJ. Nuclear γ2-positive cells were quantified as the percentage of cells with a nuclear/cytoplasmic signal ratio > 1.0. 3-5 fields of view (FOVs) were analyzed per condition per experiment. Colocalization was assessed by intensity profiles generated with the “Plot Profile” function. For Masson’s trichrome and PAS staining, positive areas were measured by “Color Deconvolution” followed by threshold analysis. TUNEL-positive and DAPI-positive nuclei were counted with “Analyze Particles”, and the percentage of TUNEL-positive cells was calculated. Immunohistochemistry staining intensity was analyzed with the IHC Profiler plugin, and H-scores were calculated as (P1 × 1) + (P2 × 2) + (P3 × 3), where P1, P2, and P3 represent the percentages of low, medium, and high positive staining, respectively. For each sample, 3-5 FOVs were analyzed.

### Statistical analysis

Data were analyzed with GraphPad Prism 9.0 and are presented as mean ± SD. Statistical tests are described in figure legends; P < 0.05 was considered significant. Experiments were repeated at least three times unless noted.

## Supplementary information

Supplementary information contains 5 figures and 3 tables.

## Acknowledgments

We thank Drs. X. Wu and T. Xu (Fudan University) for the piggyBac system, and the Medical Science Public Platform of Shenzhen Campus, Sun Yat-sen University, for imaging assistance.

## Declaration of interests

The authors declare no competing financial interests.

## Funding

This project is supported by the Shenzhen Science and Technology Program (JCYJ20240813151133043 to X. X.) and the Guangdong Basic and Applied Basic Research Foundation (2023A1515011923 to X. X; 2024A1515010663 to H.T.), and the National Natural Science Foundation of China (82470365 & 82170357 to H.T.).

## Author contributions

X. Xie and C. Li conceived the project. C. Li, T. Yi, Y. Cui, H. Tan, and X. Xie designed and performed experiments, with contributions from B. Cheng, J. Lan, C. Zhang, C. Lin, F. Yang, and Y. Chen. X. Xie and C. Li analyzed data and wrote the manuscript with input from all authors. B. Zhao, X. Wang, H. Peng, and L. Yan provided reagents and materials. X. Xie and H. Tan supervised the project and acquired funding.

## Data availability

All data necessary to evaluate the conclusions of this study are presented in the paper and/or the Supplementary Materials. The RNA-seq data have been deposited in the National Center for Biotechnology Information (NCBI) Gene Expression Omnibus (GEO) database and are accessible under GEO series accession number GSE328904.

## Abbreviations

Dox: doxorubicin
DIC: Dox-induced cardiomyopathy
AMPK: AMP-activated protein kinase
NLS: nuclear localization signal
γ2-AMPK: γ2-containing AMPK
ACC: acetyl-CoA carboxylase
RPs: ribosomal proteins
HDAC3: Histone deacetylase 3
TIP60: Tat-interactive protein 60 kDa
FBL: fibrillarin
MDM2: mouse double minute 2
Pol I: RNA polymerase I;
TIF-IA: transcription initiation factor IA

## Notes

### Competing Interest Statement

The authors have declared no competing interest.

