## Supplementary information for "AMPKγ2 Deacetylation Drives Nuclear Translocation and Doxorubicin-Induced Cardiomyopathy via Nucleolar Stress Signaling"

Contents

Supplementary Figures S1–S5 (with legends).
Supplementary Table S1. GSEA enrichment of the KEGG p53 signaling pathway
Supplementary Table S2. Antibodies used in this study.
Supplementary Table S3. DNA oligonucleotides used in this study.

### Supplementary Figure and legends

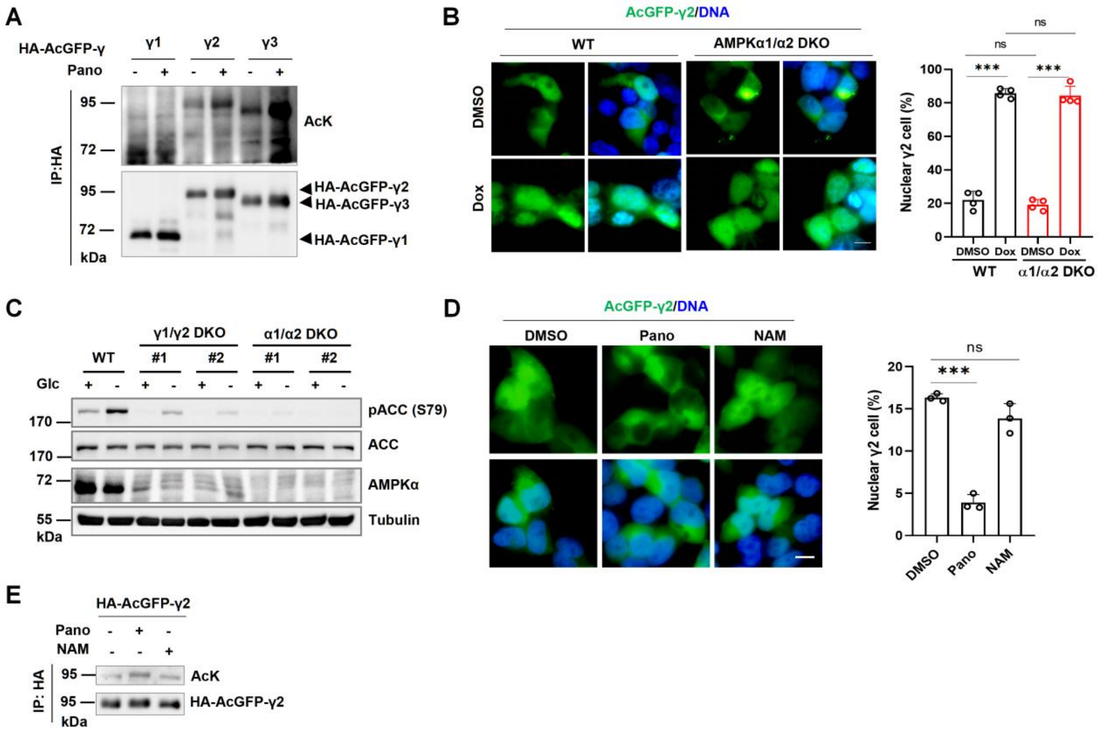


**Fig. S1. Dox-induced γ2 deacetylation and nuclear translocation are independent of Sirtuins and AMPK activity**

**(A)** IB analysis of γ subunit acetylation. 293T cells transfected with HA-AcGFP-tagged γ1, γ2, or γ3 were treated with Pano (4 μM, 12 h). γ acetylation was detected with an anti-acetyl-lysine (AcK) antibody after HA IP.

**(B)** Subcellular distribution of AcGFP-γ2 in WT and AMPKα1/α2 DKO 293T cells. Representative images (left); quantified nuclear γ2-positive cells (right). Scale bar, 10 μm.

**(C)** IB analysis of AMPK activity in AMPKγ1/γ2 DKO and AMPKα1/α2 DKO 293T cells. Cells were cultured with (+) or without (–) glucose, and AMPK activity was assessed by pACC (Ser79).

**(D)** IF analysis of γ2 subcellular distribution upon Pano or NAM treatment. 293T cells stably expressing AcGFP-γ2 were treated with DMSO, Pano (4 μM, 12 h), or NAM (2.5 mM, 12 h). Representative images (left); quantification of nuclear γ2-positive cells (right). Scale bar, 10 μm.

**(E)** IB analysis of γ2 acetylation upon Pano or NAM treatment. 293T cells stably expressing HA-AcGFP-γ2 were treated as indicated, and γ2 acetylation was detected with an AcK antibody after HA immunoprecipitation.

Data are mean ± SD. ***$P < 0.001$; ns, not significant. Statistical significance was assessed by two-tailed Student's t-test.

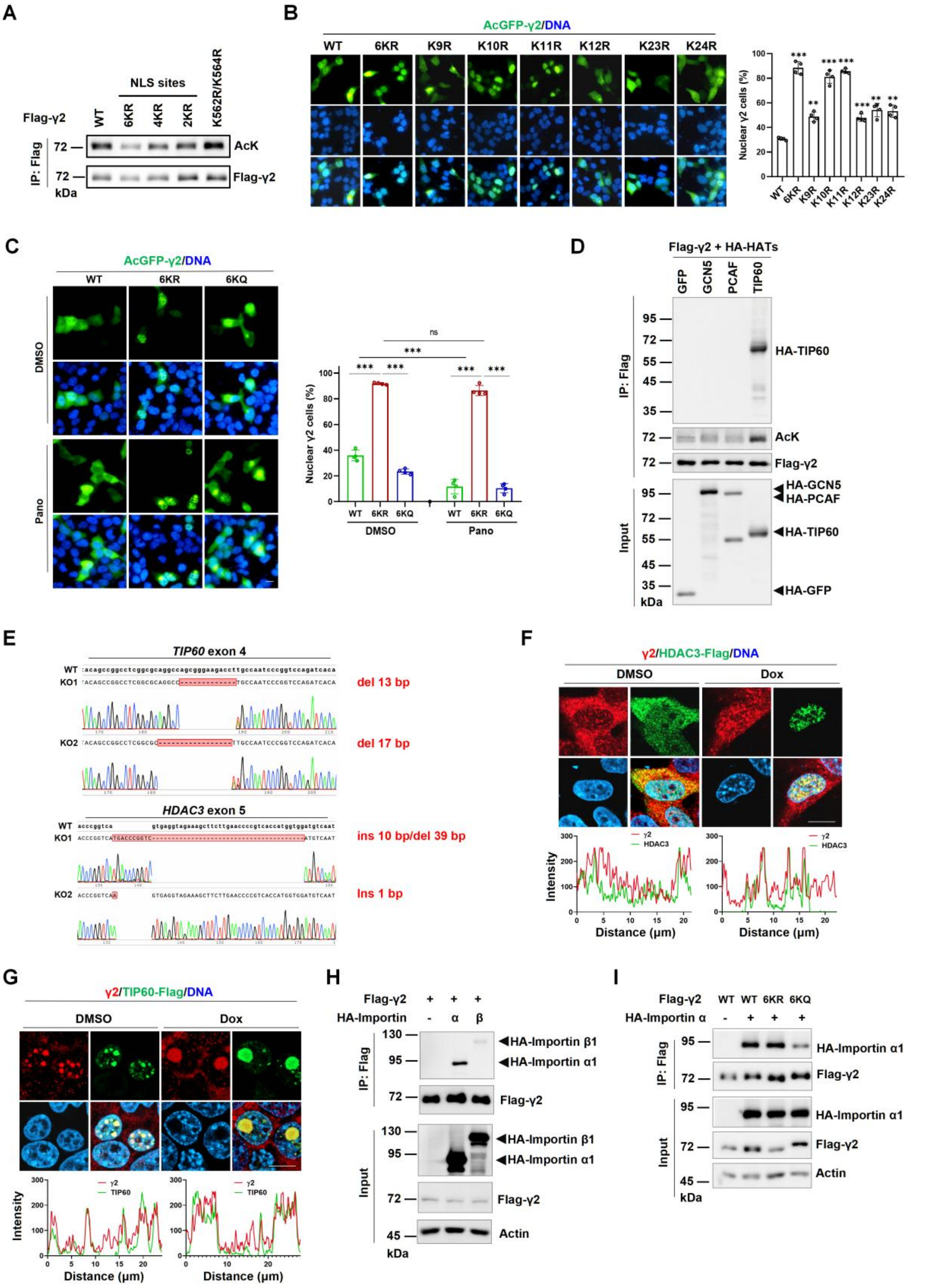


**Fig. S2. γ2 acetylation sites and regulatory mechanisms for nucleocytoplasmic shuttling**

**(A)** IB analysis of γ2 acetylation in KR mutants. 293T cells expressing Flag-γ2 WT, 6KR, 4KR, 2KR, or K56R/K564R were immunoprecipitated with anti-Flag resin, and acetylation was detected with AcK antibody.

**(B)** Subcellular distribution of single lysine mutants. 293T cells expressing AcGFP-γ2 KR mutants were analyzed by IF. Representative images (left); quantification of nuclear γ2-positive cells (right). Scale bar, 10 μm.

**(C)** Effect of Pano on nuclear accumulation of γ2 mutants. 293T cells expressing AcGFP-γ2 WT, 6KR, or 6KQ were treated with DMSO or Pano (4 μM, 12 h). Representative images (left); quantification (right). Scale bar, 10 μm.

**(D)** IP analysis of γ2 acetylation by TIP60. 293T cells were cotransfected with Flag-γ2 and indicated HA-tagged HAT constructs. Cell lysates were prepared 36 h post-transfection, and CoIP was performed using anti-Flag resin, and acetylation was detected with AcK antibody

**(E)** Sanger sequencing of CRISPR/Cas9-edited genomic loci in WT and HDAC3/TIP60 KO cell lines, del: deletion; ins: insertion.

**(F-G)** Colocalization of γ2 with HDAC3-Flag **(F)** or TIP60-Flag **(G)** upon Dox treatment. 293T cells were transfected with indicated constructs, treated with Dox (1.25 μM, 12 h), and analyzed by IF. Intensity profiles show line scan analysis of signal colocalization. Scale bar, 10 μm.

**(H-I)** CoIP analysis of γ2-importin interaction. **(H)** 293T cells were cotransfected with Flag-γ2 and HA-importin α1 or β1. **(I)** γ2 WT, 6KR, or 6KQ was cotransfected with HA-importin α1, and interactions were analyzed by CoIP using anti-Flag resin.

Data are mean ± SD. **P < 0.01, ***P < 0.001; ns, not significant. Statistical significance was assessed by two-tailed Student's t-test.

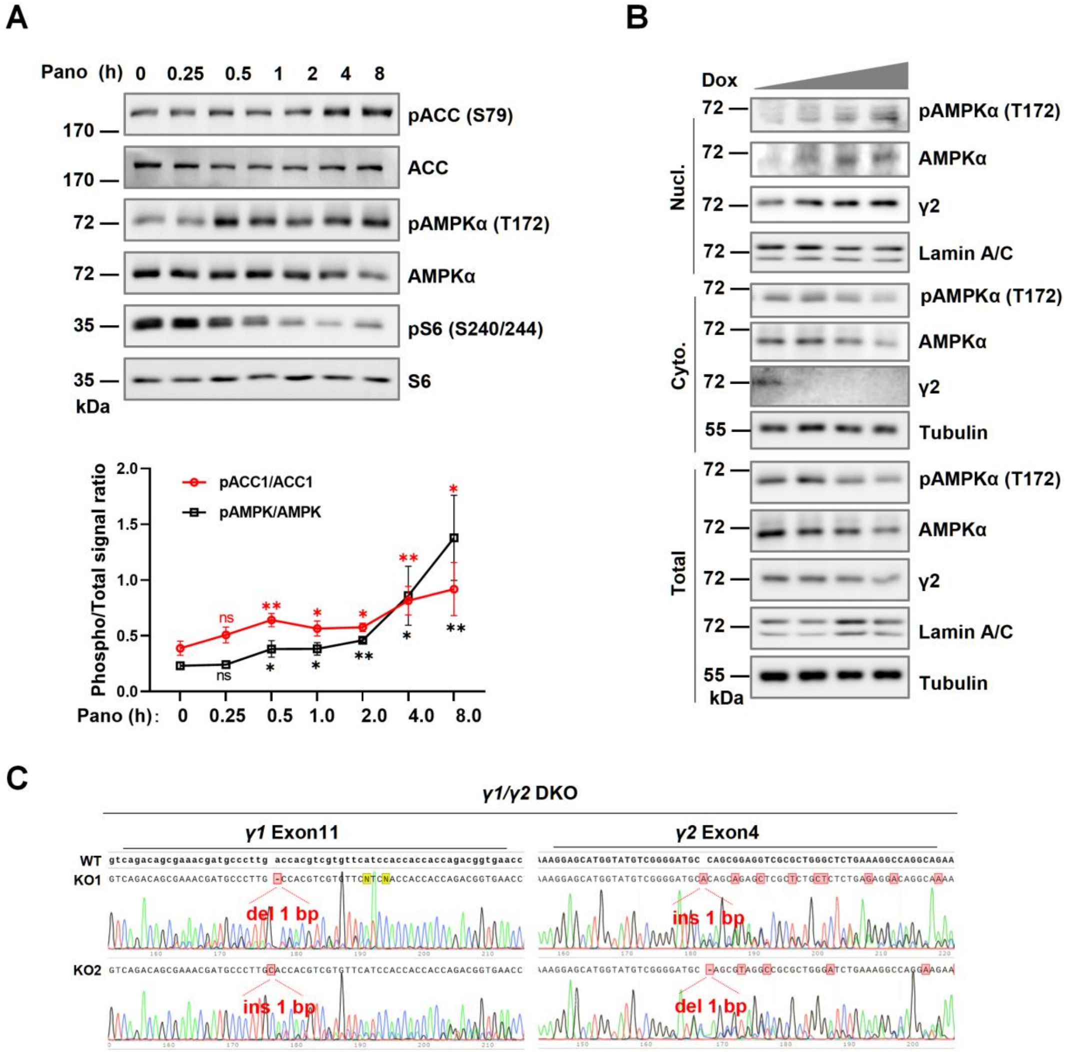


**Fig. S3. Dox differentially regulates nuclear and cytosolic AMPK activity**

**(A)** Time course of panobinostat-induced AMPK activation. 293T cells were treated with Pano (4 μM) for the indicated times. AMPK activity was assessed by pACC (Ser79), pAMPKα (Thr172), and pS6 (Ser240/244). Quantification shows phospho/total signal ratios (n = 3).

**(B)** Dose-dependent nuclear AMPKα activation by Dox. HL-1 cells were treated with escalating doses of Dox (0, 0.625, 1.25, 2.5 μM, 12 h). Subcellular fractionation was performed, and the nuclear extracts, cytosolic extracts, and total lysates were analyzed by IB for γ2 distribution and AMPKα phosphorylation.

**(C)** Sanger sequencing of CRISPR/Cas9-edited genomic loci in γ1/γ2 DKO cell lines, del: deletion; ins: insertion.

Data are mean ± SD. *P < 0.05, **P < 0.01; ns, not significant. Statistical significance was assessed by two-tailed Student's t-test.

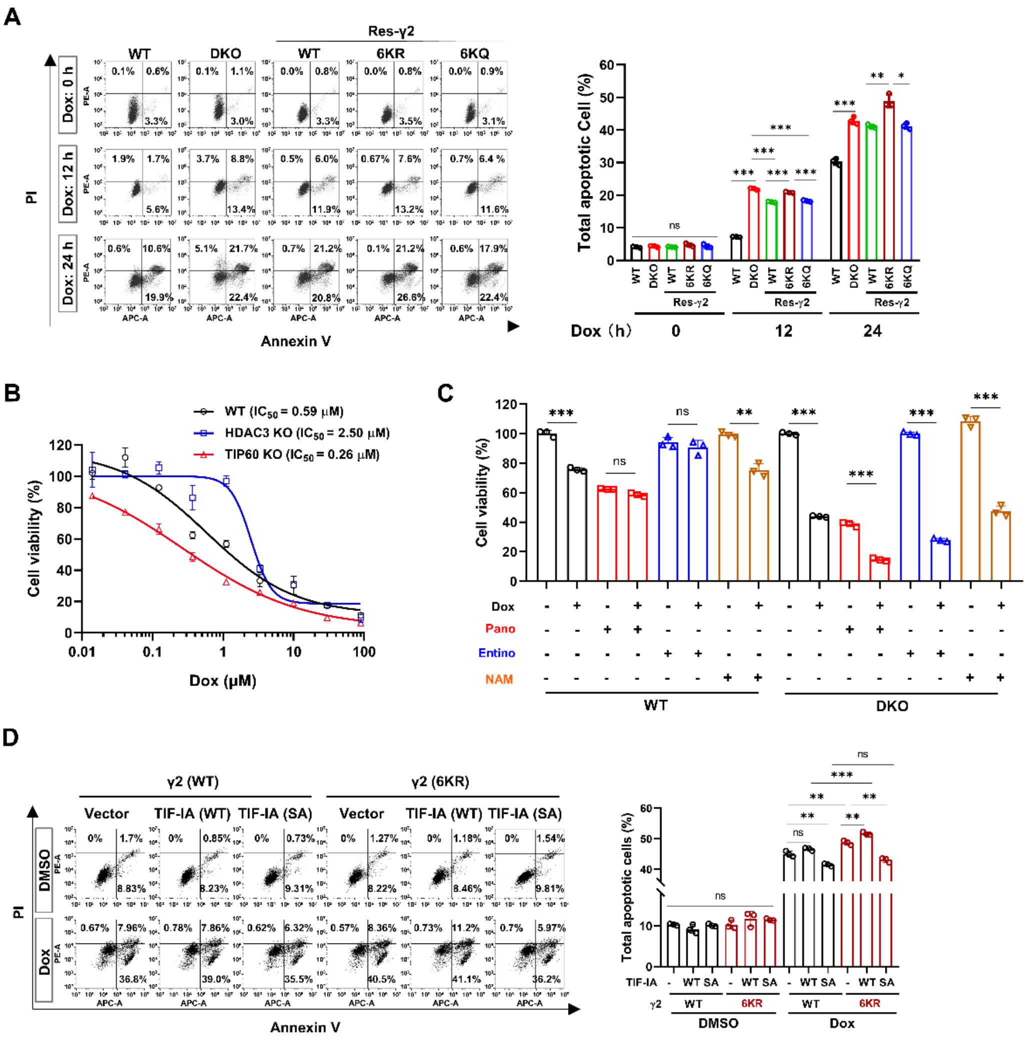


**Fig. S4. γ2 acetylation protect cardiomyocytes from Dox-induced apoptosis.**

**(A)** Flow cytometry analysis of apoptosis by Annexin V/PI staining. γ2 variant-rescued DKO HL-1 cells were treated with Dox for the indicated times. Total apoptosis was defined as the percentage of early apoptotic Annexin V$^{+}$/PI$^{-}$ cells plus late apoptotic Annexin V$^{+}$/PI$^{+}$ cells (n = 3 biological replicates).

**(B)** Dose-response curves in HDAC3 KO and TIP60 KO cells. 293T WT or KO cells were treated with escalating doses of Dox (0, 0.014, 0.04, 0.12, 0.37, 1.11, 3.33, 10, 30, 90 μM, 24 h). Cell viability was determined by CCK-8 assay. Dose-response curves were plotted and $IC_{50}$ values were fitted using GraphPad Prism 9.0.

**(C)** Cell viability upon Dox, HDACi, or co-treatment in HL-1 cells. Cells were treated with Dox (1.25 μM, 24 h), panobinostat (4 μM, 24 h), entinostat (4 μM, 24 h), or NAM (2.5 mM, 24 h) individually or in combination. Cell viability was determined by CCK-8 assay.

**(D)** TIF-IA rescues 6KR-induced apoptosis. γ2 WT- or 6KR-rescued DKO HL-1 cells expressing TIF-IA WT or S635A mutant were analyzed by Annexin V/PI staining. Total apoptosis was defined as the percentage of early apoptotic Annexin V$^{+}$/PI$^{-}$ cells plus late apoptotic Annexin V$^{+}$/PI$^{+}$cells. (n = 3 biological replicates).

Data are mean ± SD. *P < 0.05, **P < 0.01, ***P < 0.001; ns, not significant. Statistical significance was assessed by two-tailed Student's t-test.

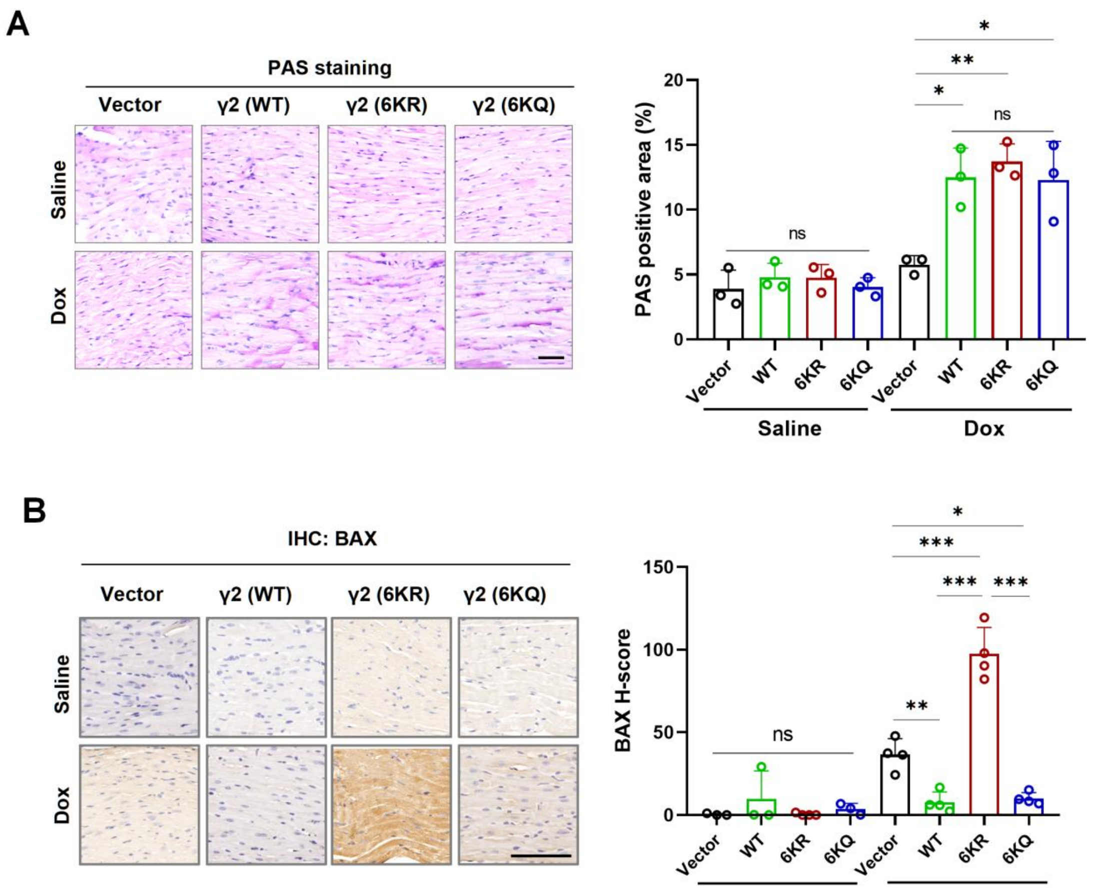


**Fig. S5. γ2 acetylation status regulates pro-apoptotic signaling but not glycogen content in DIC.**

**(A)** Glycogen staining of heart sections from DIC mice. Representative PAS staining images showing glycogen deposition (purple). PAS-positive areas were quantified using the “Color Deconvolution” function followed by threshold analysis in ImageJ (n = 3). Scale bar, 50 μm.

**(B)** Immunohistochemical staining of BAX. Representative BAX staining images (left). BAX-positive areas were quantified using IHC Profiler plugin, and H-scores were calculated (n = 4). Scale bar, 50 μm.

Data are mean ± SD. *P < 0.05, **P < 0.01, ***P < 0.001; ns, not significant. Statistical significance was assessed by two-tailed Student's t-test.

**Supplementary Table S1. GSEA of the KEGG p53 signaling pathway**

| KEGG p53 signaling pathway | | | |
|---|---|---|---|
| **Group** | **NES** | **Adjusted p-value** | **Core enrichment** |
| 6KR vs WT | 1.703 | 0.0045 | Cdkn1a, Zfp385a, Ppp1r13l, Mdm2, Gtse1, Cd82, Rrm2, Shisa5, Apaf1, Fas, Pidd1, Aifm2, Steap3, Ccnb1, Sesn2, Sesn1, Tnfrsf10b, Zmat3, Ppm1d, Ei24, Tsc2, Ccnb2, Cdk1 |
| 6KQ vs WT | -1.265 | 0.1098 | Siah1b, Adgrb1, Gadd45a, Ccnd2, Ccnd1, Gadd45b, Bax, Siva1, Ccne1, Gadd45g, Cdk2, Bcl2l1, Bid, Trp53, Cdk4, Thbs1 |
| 6KR vs 6KQ | 1.885 | 0.0001 | Cdkn1a, Rrm2, Trp53, Zfp385a, Gtse1, Ppp1r13l, Shisa5, Bax, Cdk4, Cdk1, Sesn2, Aifm2, Ccnd1, Siva1, Ccnd3, Ei24, Cd82, Ccnb2, Sfn, Bbc3, Steap3, Cdk2, Pidd1, Tsc2, Tnfrsf10b |

**Supplementary Table S2. Antibodies used in this study**

| Antibody Name | Dilution | Vendor (Catalog #) |
|---|---|---|
| Anti-Acetylated-lysine | 1:1000 | Cell Signaling Technology (#9441S) |
| Anti-pAMPKα (Thr172) | 1:1000 | Cell Signaling Technology (#2535S) |
| Anti-Lamin A/C | 1:1000 | Cell Signaling Technology (#4777S) |
| Anti-AMPKγ2 | 1:2000 | Sigma (#HAP004246) |
| Anti-AMPKγ1 | 1:1000 | HUABIO (#HA722691) |
| Anti-AMPKα | 1:1000 | HUABIO (#ET1608-40) |
| Anti-pACC (S79) | 1:1000 | HUABIO (#HA721714) |
| Anti-ACC1 | 1:1000 | HUABIO (#ET1609-77) |

| | | |
|---|---|---|
| Anti-HDAC3 | 1:2000 | HUABIO (#ET1610-5) |
| Anti-p53 | 1:2000 | HUABIO (#HA601315) |
| Anti-cleaved/pro Caspase-3 | 1:1000 | HUABIO (#ET1608-64) |
| Anti-HA | 1:20000 | HUABIO (#HA721750) |
| Anti-Flag | 1:5000 | HUABIO (#HA601167) |
| Anti-HA magnetic beads | - | HUABIO (#HAK21042) |
| Anti-Flag magnetic beads | - | HUABIO (#HAK21011) |
| Anti-RPS6 | 1:1000 | ABclonal (#A11874) |
| Anti-pS6 (S240/S244) | 1:1000 | Solarbio (#K006232P) |
| Anti-GAPDH (3B3) | 1:5000 | Abmart (#M20006) |
| Anti-β-Tubulin (C66) | 1:5000 | Abmart (#M20005) |
| Anti-β-Actin | 1:5000 | Abmart (#M40104) |
| Anti-pan-phospho-ser/thr/tyr | 1:1000 | Abmart (#M210030) |
| Anti-FBL | 1:300 (IF) | proteintech (#66985-1-Ig) |
| Anti-RPL11 | 1:2000 | proteintech (#16277-1-AP) |
| HRP-conjugated Rabbit Anti-Mouse IgG | 1:10000 | proteintech (#SA00001-19) |
| HRP-conjugated Mouse Anti-Rabbit IgG | 1:10000 | proteintech (#SA00001-7L) |
| HRP-labeled goat anti-mouse IgG (H+L) | 1:3000 | Beyotime (#A0216) |
| HRP-labeled goat anti-rabbit IgG (H+L) | 1:3000 | Beyotime (#A0208) |
| Goat anti-mouse IgG (Alexa Fluor® 647) | 1:1000 | Abcam (#ab150115) |
| Goat anti-rabbit IgG (Alexa Fluor® 488) | 1:1000 | Abcam (#ab150077) |
| Goat anti-mouse IgG (Alexa Fluor® 488) | 1:1000 | Abcam (#ab175660) |
| Goat anti-rabbit IgG (Alexa Fluor® 647) | 1:1000 | Abcam (#ab150079) |

**Supplementary Table S3. DNA oligonucleotides used in this study.**

| | | |
|---|---|---|
| **Primers for RT–qPCR** | | |
| Gene | Primer | Sequence (5'-3') |
| 45S pre-rRNA (ITS1) | Forward | TCTCGTTTCGTTCCTGCTGG |
| | Reverse | GATCCACCGCTAAGAGTCGTATC |
| Ubc | Forward | GAGCCCAGTGTTACCACCAAGAAG |
| | Reverse | CACACCCAAGAACAAGCACAAGGAG |
| **sgRNA sequences** | | |
| sgRNA | Primer | Sequence (5'-3') |
| sgTIP60 #1 (human) | Forward | CACCGGATTGGCAAGGTCTTCCCGC |
| | Reverse | AAACGCGGGAAGACCTTGCCAATCC |
| sgHDAC3 #1 (human) | Forward | CACCGCCGTCATGACCCGGTCAGTG |
| | Reverse | AAACCACTGACCGGGTCATGACGGC |
| sgAmpkγ1 (Mouse) | Forward | CACCGGATGAACACGACGTGGTCAA |
| | Reverse | AAACTTGACCACGTCGTGTTCATCC |
| sgAmpkγ2 (Mouse) | Forward | CACCGTGGTATGTCGGGGATGCCAG |
| | Reverse | AAACCTGGCATCCCCGACATACCAC |
| **Amplicon primers for Cas9/sgRNA target region** | | |
| Gene | Primer | Sequence (5'-3') |
| TIP60 (human) | 5' out | TCCCTCTCACCCTGACTTCAT |
| | 3' out | TCCCTGGAATTGCTGAGGGT |
| HDAC3 (human) | 5' out | GAGGAGGGGACACCTGAGAT |
| | 3' out | TCACAATTCGAGACCCGGTG |
| Ampkγ1 (Mouse) | 5' out | TGGAGCTCCTGAGAGAAGACT |
| | 3' out | GTCAGGTCGAGGCAGTTACC |
| Ampkγ2 (mouse) | 5' out | GTTGCTCTCAGCCTGTGACT |
| | 3' out | AACAGTCAGGGCTGTATGCC |
